# Evaluating primer and probe mismatch tolerance in an Influenza A matrix gene RT-qPCR using contemporary human and zoonotic strains

**DOI:** 10.64898/2026.02.23.707407

**Authors:** Caroline Amuche Okoli

## Abstract

**Background:** Genetic drift and host-associated adaptation in influenza A viruses threaten the long-term reliability of RT-qPCR–based diagnostics, particularly when nucleotide mismatches arise within primer and probe binding regions. Conventional assay evaluations often emphasize sequence conservation but rarely assess functional mismatch tolerance across divergent subtypes and hosts.

**Methods:** We performed an in silico evaluation of a matrix (M) gene–targeted RT-qPCR assay by aligning primer and probe binding regions against 22 H1N1 isolates and representative H3N2 and H5N1 reference strains, including recent zoonotic isolates from avian and bovine hosts. Nucleotide mismatches were identified, quantified, and mapped relative to assay components and oligonucleotide termini. Mismatch burden was summarized by subtype and assay region.

**Results:** H1N1 isolates exhibited complete conservation across primer and probe regions. In contrast, H3N2 and H5N1 strains demonstrated subtype-specific sequence variability, with a total of eleven mismatches identified across seven non-H1N1 isolates (mean mismatch per isolate = 2.43). Probe mismatches predominated (63.6%), occurring primarily at internal positions, while primer mismatches were infrequent and largely avoided 3′ terminal nucleotides. Recent H5N1 isolates (2023–2024) shared conserved internal mismatches in the probe and forward primer, whereas a historical H5N1 isolate (2016) exhibited a distinct profile including a terminal probe mismatch. Despite this variability, mismatch patterns were consistent with preserved amplification potential.

**Conclusion:** This study demonstrates that the evaluated influenza A M gene RT-qPCR assay exhibits inherent mismatch tolerance across human and zoonotic subtypes. By shifting diagnostic evaluation from strict sequence identity to functional resilience, our findings provide a framework for designing and maintaining robust molecular assays suitable for surveillance and pandemic preparedness amid ongoing viral evolution.

**Graphical Abstract:** In silico evaluation of an influenza A matrix gene RT-qPCR assay demonstrates subtype-specific primer and probe mismatches across H3N2 and H5N1 strains, including recent zoonotic isolates. Despite observed variability, mismatches predominantly occur at internal positions and spare primer 3′ termini, supporting inherent assay mismatch tolerance and suitability for surveillance applications.

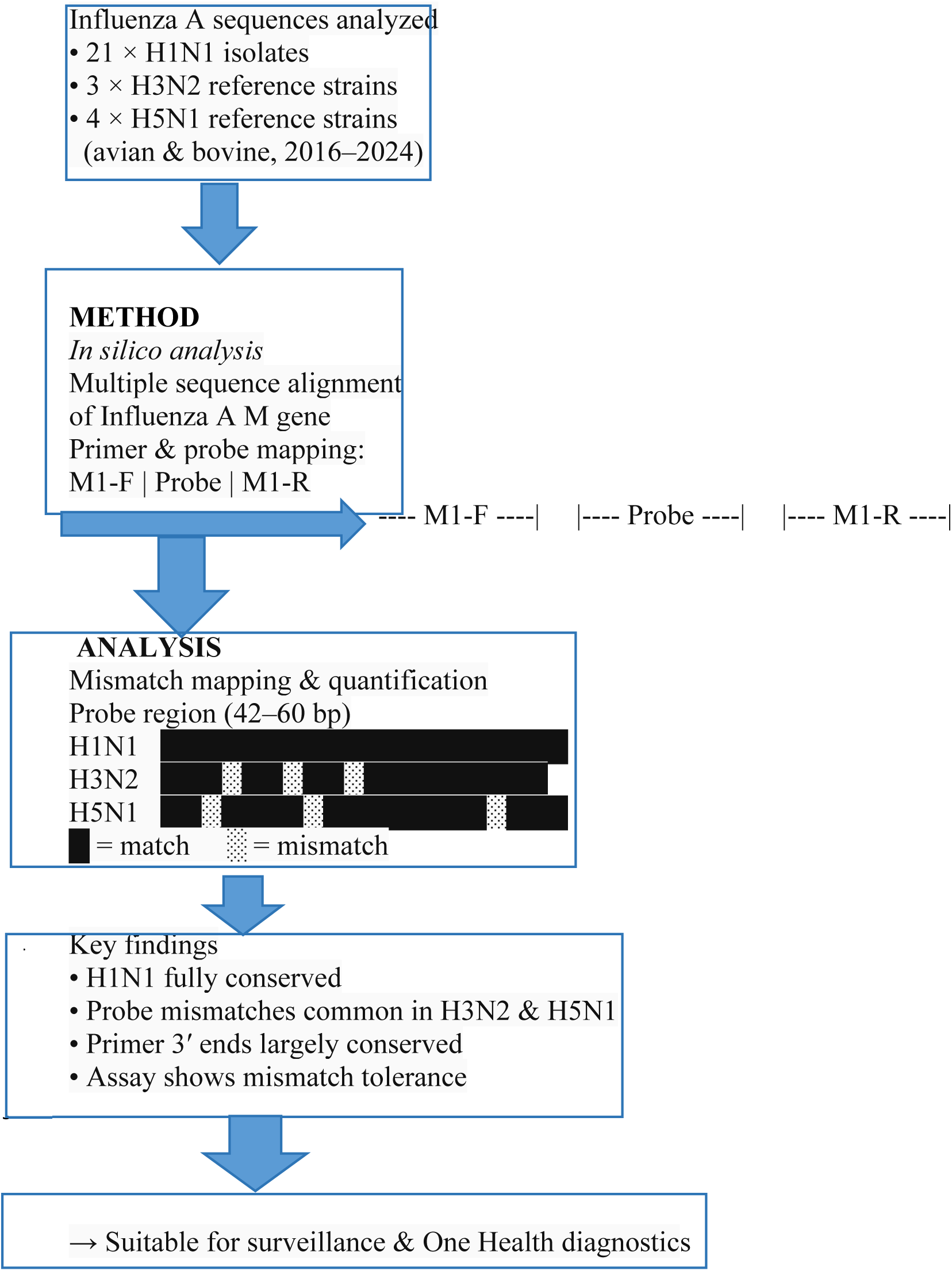

## Introduction

Influenza A viruses remain a persistent global public health threat, causing annual seasonal epidemics and posing continual pandemic risk through zoonotic transmission and viral reassortment [1]. The wide host range of influenza A viruses including humans, swine, avian species, and other mammals facilitates ongoing genetic diversification driven by mutation, reassortment, and host adaptation. This genetic plasticity complicates disease surveillance and challenges the long-term reliability of molecular diagnostic assays used for detection and monitoring of circulating strains [2].

Real-time reverse transcription polymerase chain reaction (RT-qPCR) is the gold standard for influenza A virus detection due to its high sensitivity, specificity, and rapid turnaround time [3–4]. Most widely deployed assays target conserved genomic regions, particularly the matrix (M) gene, which encodes structural proteins essential for viral assembly and replication. The M gene is therefore favored for broad influenza A detection across subtypes and hosts and forms the basis of many World Health Organization (WHO) and Centers for Disease Control and Prevention (CDC)-recommended diagnostic protocols [3–4].

Despite targeting conserved regions, RT-qPCR assays are inherently vulnerable to nucleotide mismatches within primer and probe binding sites. Even single nucleotide substitutions can alter oligonucleotide hybridization efficiency, affect polymerase extension, or disrupt probe cleavage, potentially leading to reduced assay sensitivity or false-negative results[5]. The impact of mismatches depends on their number, position, and nucleotide context, with primer 3′ terminal mismatches generally exerting the greatest effect, while internal probe mismatches may have more modest consequences [6]. However, systematic evaluation of mismatch tolerance is rarely incorporated into routine assay design or validation workflows.

The challenge posed by sequence variability is particularly relevant in the context of influenza A virus surveillance at the human-animal interface [7]. Highly pathogenic avian influenza viruses, such as H5N1, have demonstrated increasing geographic spread and expanding host range, including recent detections in mammalian species. Concurrently, seasonal human subtypes such as H3N2 continue to evolve under immune selection pressure. These dynamics raise concerns regarding the ability of existing molecular assays to maintain performance across divergent and emerging lineages, especially when assays are extrapolated beyond their originally intended target subtypes.

Traditional diagnostic assay evaluation often focuses on analytical sensitivity, specificity, and limit of detection using a limited set of reference strains [8]. While these metrics are essential, they do not fully capture assay robustness under real-world conditions of viral evolution. In silico approaches that explicitly map and quantify primer and probe mismatches across diverse and contemporary viral sequences provide a complementary strategy to anticipate diagnostic vulnerabilities before clinical failure occurs. Such analyses are increasingly recognized as valuable tools for proactive assay assessment, particularly when experimental validation materials are limited or emerging strains are difficult to access.

In this study, we performed a comprehensive in silico evaluation of a matrix gene–targeted influenza A RT-qPCR assay to assess primer and probe mismatch patterns across multiple subtypes and hosts. Using a dataset comprising 22 H1N1 isolates alongside representative H3N2 and H5N1 reference strains including recent zoonotic isolates, we mapped nucleotide variability within primer and probe binding regions, quantified mismatch burden, and examined positional distribution relative to oligonucleotide termini. By focusing on mismatch tolerance rather than strict sequence identity, this work aims to provide a framework for evaluating diagnostic resilience in the face of ongoing influenza A virus evolution and to inform the design of robust molecular assays suitable for long-term surveillance and pandemic preparedness.

## Materials and Methods

### Sequence Retrieval

Influenza A virus matrix (M) gene sequences were retrieved from curated public databases (NCBI) to represent genetic diversity across host species, geographic locations, and years of isolation. The dataset comprised 22 H1N1 isolates and reference sequences representing H1N1, H3N2, and H5N1 subtypes. Selected sequences included isolates from swine, canine, avian, bovine, and other mammalian hosts collected across multiple regions and time periods.

Accession numbers, host species, country of origin, and year of submission for all sequences included in the analysis are provided in Table 1.

**Table 1:** Metadata of the samples.

| Subtype | Accession/isolates | Location | Isolation source | Host | Year of collection | Year of submission |
| --- | --- | --- | --- | --- | --- | --- |
| H1N1 | ON822118.1 | Guatemala | Nasal swab | Swine | 2016 | 2022 |
|  | ON822150.1 |  |  |  |  |  |
|  | ON822156.1 |  |  |  |  |  |
|  | ON822162.1 |  |  |  |  |  |
|  | ON822132.1 |  |  |  |  |  |
|  | ON822137.1 |  |  |  |  |  |
|  | ON822156.1 |  |  |  |  |  |
|  | ON822144.1 |  |  |  |  |  |
|  | ON822119.1 |  |  |  |  |  |
|  | ON822302.1 |  |  |  |  |  |
|  | ON822309.1 |  |  |  |  |  |
|  | ON822315.1 |  |  |  |  |  |
|  | ON822325.1 |  |  |  |  |  |
|  | ON822322.1 |  |  |  |  |  |
|  | ON822125.1 |  |  |  |  |  |
|  | ON823112.1 |  |  |  |  |  |
|  | ON823120.1 |  |  |  |  |  |
|  | ON823128.1 |  |  |  |  |  |
|  | ON823136.1 |  |  |  |  |  |
|  | MK680249.1 | India (Mizoram) | Nasal swab | Swine | 2017 | 2019 |
|  | LC662543.1 | Japan | MDCK cell line | Canis lupus familiaris | 2021 | 2021 |
|  | NC_002016 | Puerto Rico | - | Human | 1934 | 2000 |
| H3N2 | KJ609209.1 | Australia | - | Human | 2009 | 2014 |
|  | NC_007367.1 | Tompkins county (USA) | Boy | Human | 2004 | 2005 |
|  | PQ731530.1 | California (USA) | Clinical | Human | 2024 | 2024 |
| H5N1 | NC_007363.1 | Guangdong (Hong Kong) | - | Goose | 1996 | 1999 & 2005 |
|  | OR819343.1 | New york (USA) | Swab | Chicken | 2023 | 2023 |
|  | PP599468.1 | Texas (USA) | Milk | Bovine | 2024 | 2024 |
|  | KY614875.1 | West Java (Indonesia) | Tissue | Chicken | 2016 | 2017 |

### Multiple Sequence Alignment and Consensus Generation

All sequences were aligned using MAFFT as implemented in Geneious Prime (Biomatters Ltd.). Alignments were visually inspected to assess sequence conservation, gap distribution, and the presence of single-nucleotide polymorphisms (SNPs). No insertions or deletions were observed within the primer or probe binding regions.

A consensus sequence was generated from the alignment by selecting the most frequently occurring nucleotide at each position across the majority of sequences. This consensus sequence served as the template for primer and probe design.

### Primer and Probe Design and Characteristics

Primers and a hydrolysis probe targeting a conserved region of the influenza A M1 gene were designed based on the consensus sequence. The forward primer (M1-F) and reverse primer (M1-R) were designed to amplify a 97 bp fragment. Primer binding sites were selected from regions exhibiting complete conservation across all H1N1 sequences and minimal variability across non-H1N1 subtypes.

The hydrolysis probe was designed to anneal internally within the amplified fragment, avoiding overlap with primer binding sites. Probe placement prioritized internal positioning and balanced GC content. Where single-nucleotide variation was observed within the probe-binding region among non-H1N1 subtypes, a consensus probe sequence was selected to maximize coverage across isolates.

Primer and probe parameters; including length, GC content, melting temperature (Tm), and secondary structure potential were evaluated using Primer-BLAST and standard thermodynamic guidelines. Conserved regions were assessed both visually and computationally to ensure optimal assay design.

### Mismatch Identification and Positional Mapping

Primer and probe mismatches were identified by comparing oligonucleotide sequences against aligned influenza A sequences. Mismatches were enumerated per sequence and mapped according to their position within each oligonucleotide.

Probe mismatches were further classified based on positional location as:

**5′ terminal** (positions 1–3), **central** (positions 7–13), or **3′ terminal** (positions 17–19).

Primer mismatches were evaluated relative to the 5′ and 3′ termini, with particular attention to the 3′ terminal nucleotides due to their importance in polymerase extension.

### Mismatch Quantification

Mismatch burden was quantified by aligning primer and probe binding regions against 28 influenza A virus sequences representing H1N1, H3N2, and H5N1 subtypes. The total number of mismatches per sequence was recorded, and subtype-specific mismatch distributions were summarized. Mean mismatch values were calculated per subtype and assay component (forward primer, probe, and reverse primer).

### Estimated Ct Impact of Probe Mismatches

The potential impact of probe mismatches on cycle threshold (Ct) values was estimated based on previously published real-time PCR performance data [9]. Expected effects were categorized as follows:

**Perfect match:** baseline Ct **One internal probe mismatch:** +0.3 to +1.0 Ct

**Two internal probe mismatches:** +0.8 to +1.5 Ct

**Three or more internal probe mismatches:** potential loss of assay sensitivity

### In silico Specificity Analysis Using Primer-BLAST

Primer specificity was evaluated using NCBI Primer-BLAST against the nucleotide collection (nt) database. Analyses were performed with no organism restriction and with organism-specific restrictions to *Homo sapiens*, *Sus scrofa*, and *Canis lupus familiaris* to assess potential off-target amplification in host genomes.

Additional Primer-BLAST analyses were conducted with organism restrictions applied to Influenza A virus and individual subtypes (H1N1, H3N2, and H5N1) to evaluate subtype coverage and predicted amplicon size.

### Assessment of Sequence Variability

Sequence variability within primer and probe binding regions was assessed across all aligned sequences. Variations were documented and evaluated based on their position within primers (5′ vs 3′ termini) and within the probe (internal vs terminal positions) to assess their potential impact on assay performance.

### Assessment of Primer–Dimer Formation

Potential primer–dimer and hairpin formations were assessed in silico using Primer-BLAST, which calculates self-complementarity and 3′ self-complementarity scores based on Primer3 algorithms. Primer pairs exhibiting low self-complementarity (<5) and minimal 3′ complementarity (≤2) were considered to have low predicted dimerization risk.

## Results

An in silico workflow was employed for the design and evaluation of primers and a hydrolysis probe targeting the Influenza A virus matrix (M1) gene (Figure 1). The workflow comprised sequence retrieval, multiple sequence alignment of reference strains and isolates, identification of conserved regions, consensus sequence generation, primer and probe design, assessment of oligonucleotide thermodynamic properties, evaluation of primer–dimer potential, and in silico specificity analysis using Primer-BLAST. This approach enabled systematic assessment of sequence conservation and mismatch distribution across influenza A subtypes.

**Figure 1.**
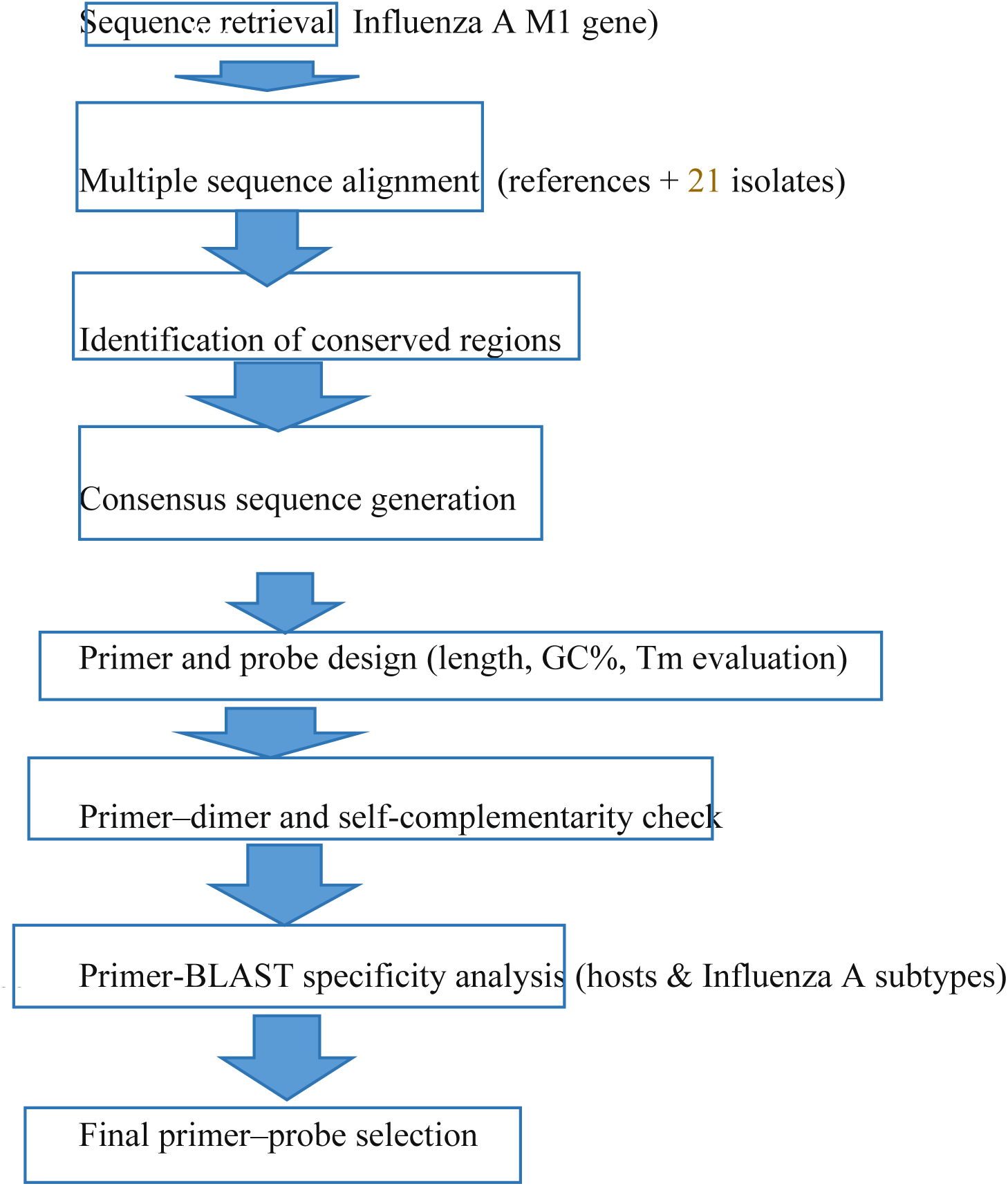
In-silico workflow used for the design and validation of primers and probe targeting the Influenza A virus M1 gene. Conserved regions were identified through multiple sequence alignment, followed by consensus-based primer and probe design and specificity assessment using Primer-BLAST.

### Sequence retrieval and dataset composition

A total of 28 Influenza A virus M1 gene sequences were analyzed, including 22 H1N1 isolates, three H3N2 reference strains, and four H5N1 reference strains (Table 1). The dataset included contemporary zoonotic H5N1 isolates from avian and bovine hosts (2023–2024) as well as an older avian H5N1 isolate from 2016, enabling assessment of both temporal and subtype-associated sequence variation.

### Multiple sequence alignment and consensus generation

Multiple sequence alignment of the M1 gene revealed strong overall conservation across all sequences analyzed, with no insertions or deletions detected within the primer or probe binding regions (Figure 2). Sequence variability was limited to single-nucleotide substitutions.

**Figure 2:**
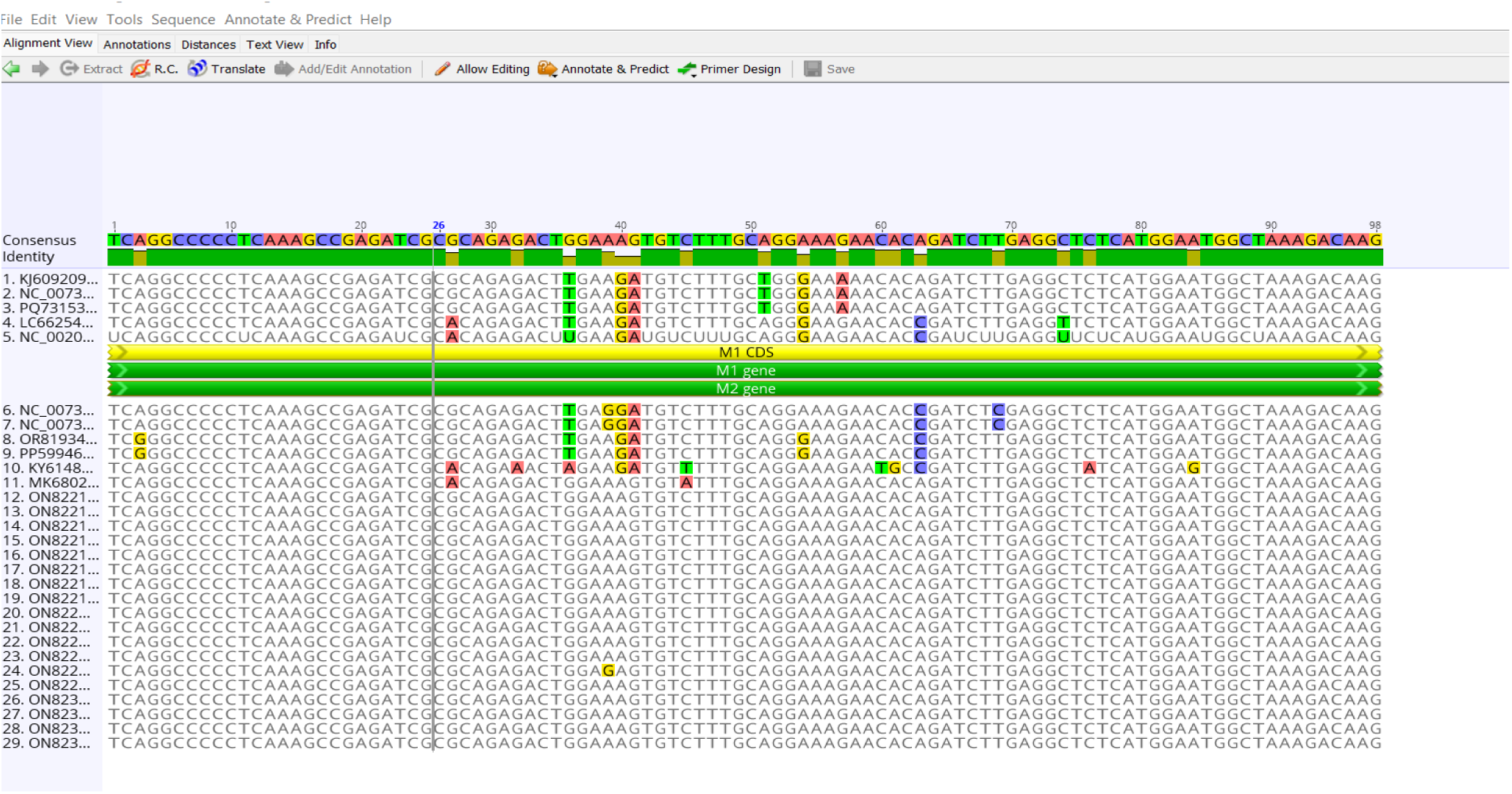
Multiple sequence alignment of the Influenza A M gene showing primer (M1-F, M1-R) and probe binding regions across H1N1, H3N2, and H5N1 sequences. Visual inspection of the alignment revealed minimal sequence variability. Nucleotide substitutions are highlighted.

**Figure 2a:**
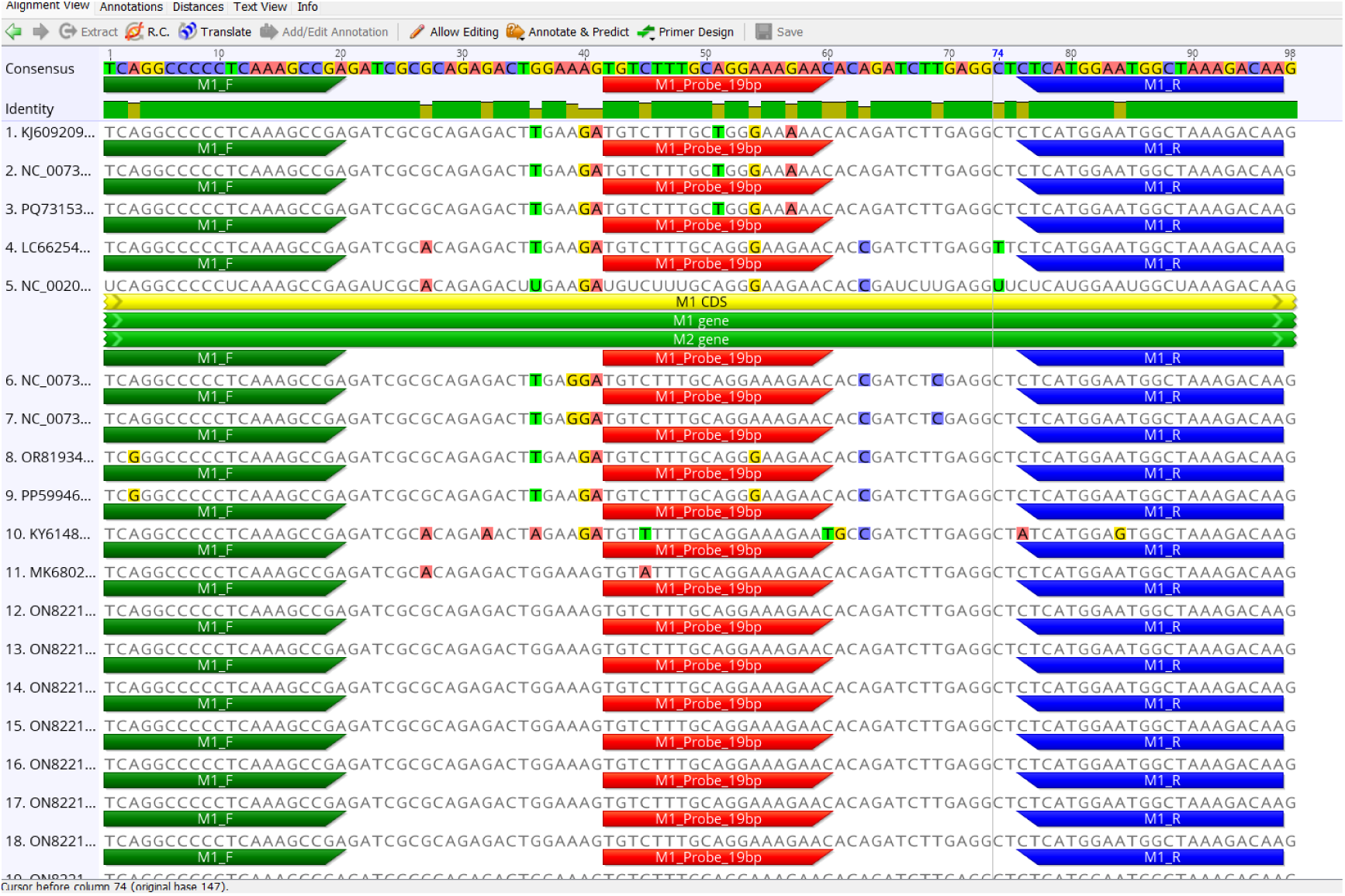
Schematic representation of the designed real-time PCR assay showing the relative positions and orientations of the forward primer, reverse primer, and hydrolysis probe within the target M1 gene region of isolates 1 to 18.Nucleotide substitutions within primer and probe regions are highlighted.

**Figure 2b:**
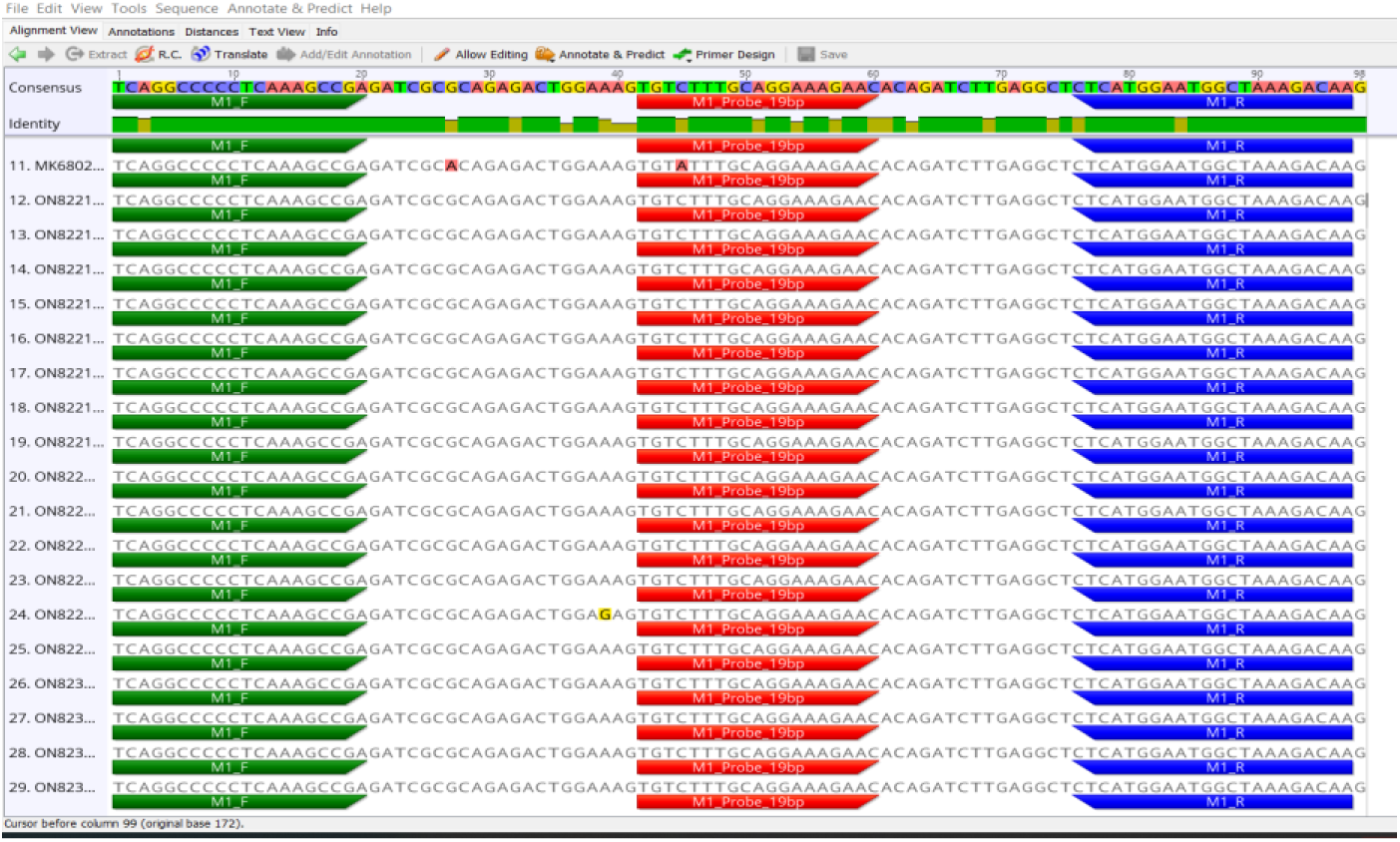
Schematic representation of the designed real-time PCR assay showing the relative positions and orientations of the forward primer, reverse primer, and hydrolysis probe within the target M1 gene region from isolates 11 to 29. Nucleotide substitutions within primer and probe regions are highlighted.

All 22 H1N1 sequences showed complete conservation across the forward primer (M1-F), probe, and reverse primer (M1-R) binding regions. In contrast, consistent subtype-specific divergence was observed in H3N2 and H5N1 sequences relative to the H1N1 consensus, with variability predominantly localized to the probe-binding region (Figure 2a–b).

## Combined subtype comparison

When all subtypes were aligned together, primer binding regions remained highly conserved across H1N1, H3N2, and H5N1 sequences, with variability confined to non-critical primer positions. Probe mismatches were subtype-specific but consistent within each lineage, suggesting structured evolutionary divergence rather than random mutation.

### Primer and Probe Design and Characteristics

Based on the consensus sequence derived from the alignment, a primer–probe set targeting a ∼97 bp region of the M1 gene was designed (Table 2). The forward primer (M1-F) was 19 bp in length with a GC content of 68.4% and a predicted melting temperature (Tm) of 64.6 °C. The reverse primer (M1-R) was 22 bp long with a GC content of 40.9% and a predicted Tm of 56.4 °C. The hydrolysis probe (M1-Probe) was 19 bp in length with an estimated GC content of ∼47% and a predicted Tm of approximately 60–62 °C.

**Table 2:**
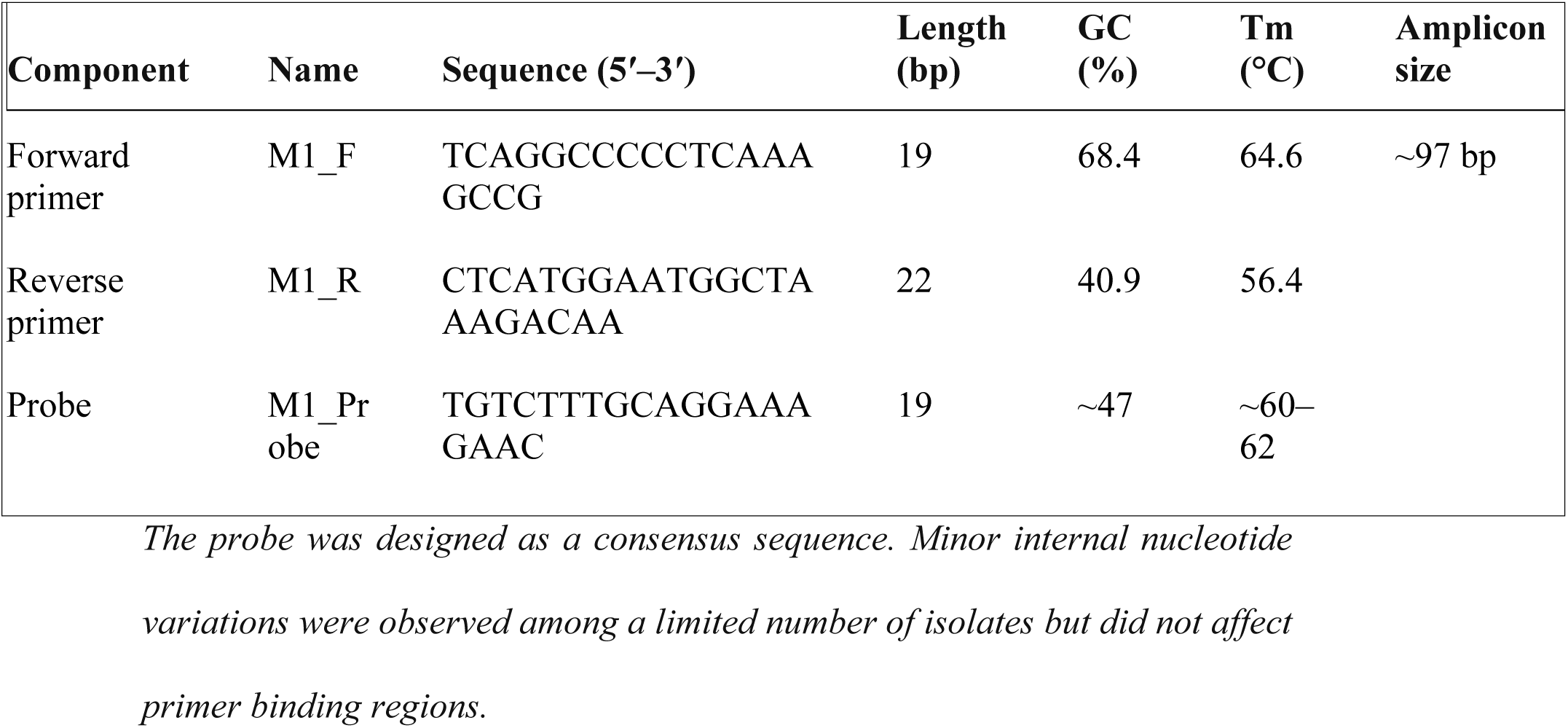
Designed primers and probe targeting the Influenza A virus M1 gene.

The probe was designed as a consensus sequence. Minor internal nucleotide substitutions were observed among a subset of non-H1N1 sequences but did not overlap with primer 3′ terminal regions.

### Primer and probe variability across subtypes

#### Primer binding regions

Primer binding regions were fully conserved across all H1N1 and H3N2 sequences analyzed. Limited primer mismatches were observed among H5N1 sequences. Two recent H5N1 isolates (OR819343.1, 2023, chicken; and PP599468.1, 2024, bovine) shared an identical A→G substitution within the forward primer binding region, while the reverse primer binding region remained conserved.

A single historical H5N1 isolate (KY614875.1, 2016, chicken) exhibited variability within the reverse primer binding region, including a C→A substitution at the 5′ terminal nucleotide and an internal A→G substitution. No primer mismatches were detected at the 3′ terminal positions of either primer across any sequence analyzed (Tables 3).

### Probe binding region

Probe variability was observed across H3N2 and H5N1 subtypes. All three H3N2 sequences exhibited identical internal probe mismatches at positions 51 bp (A→T), 54 bp (A→G), and 57 bp (G→A) relative to the H1N1 consensus sequence. No primer mismatches were observed in H3N2 isolates. Thus, each H3N2 isolate contained three probe mismatches.

Among H5N1 sequences, mismatches patterens were more heterogenous. Two recent isolates (OR819343.1 and PP599468.1) exhibited two mismatches each, comprising one forward primer mismatch and one internal probe mismatch. A historical isolate (KY614875.1) exhibited four mismatches, including two probe mismatches-one of which occurred at the probe 3’ terminal nucleotide - and two reverse primer mismatches. One H5N1 isolate showed no mismatches.

### Mismatch Positional Mapping

Mismatches were enumerated per sequence and mapped according to their position within each oligonucleotide. Probe mismatches were further classified based on positional location as: **5′ terminal** (positions 1–3), **central** (positions 7–13), or **3′ terminal** (positions 17–19). Primer mismatches were evaluated relative to the 5′ and 3′ termini, with particular attention to the 3′ terminal nucleotides due to their importance in polymerase extension (Table 3b).

**Table 3.**
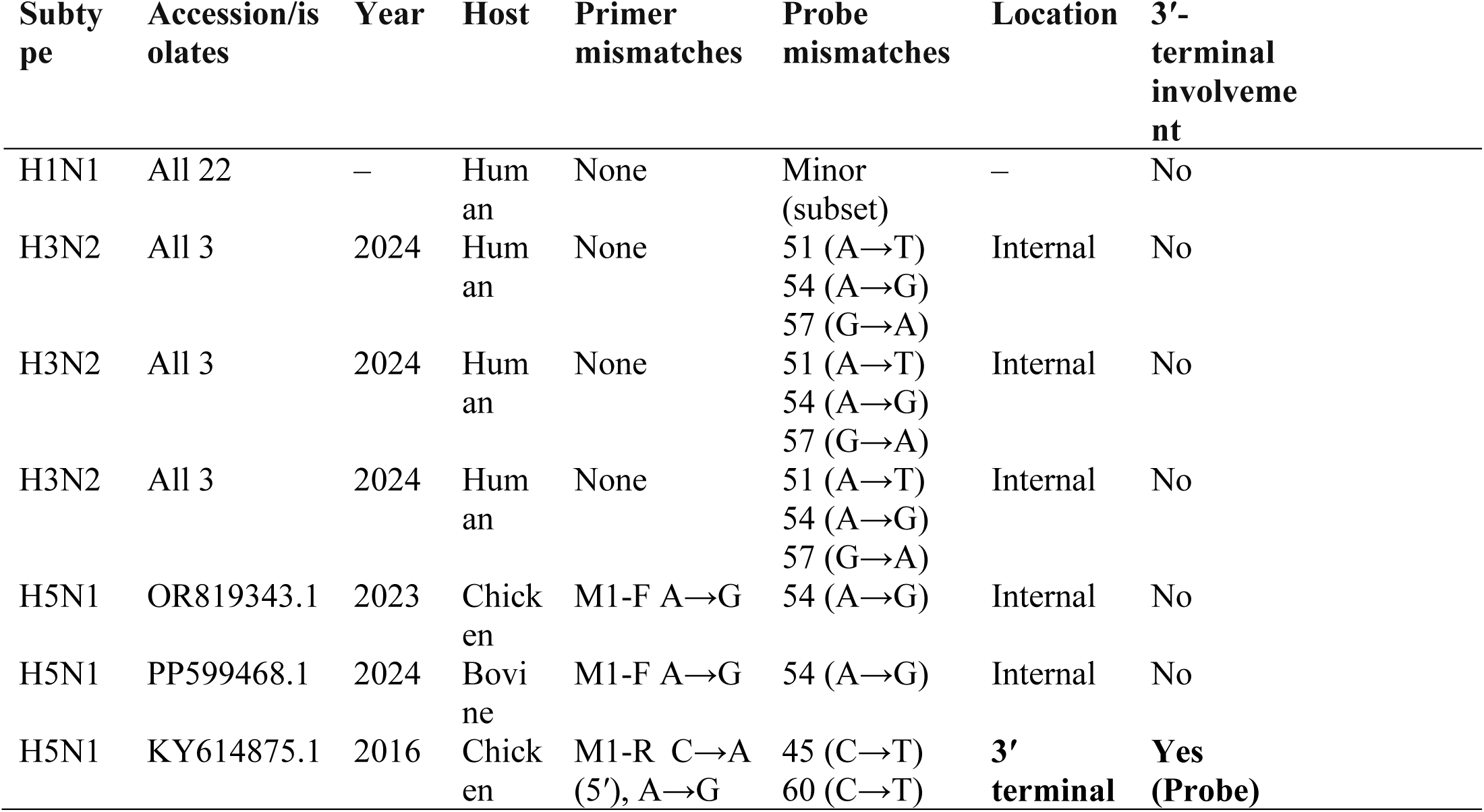
Summary of primer and probe mismatches across Influenza A subtypes.

**Table 3b.** Positional weighing of mismatches (Weight mismatches by position)

| Subtype |  |  |  |  | Position<br>s |  |  |  |  |  |  |  |
| --- | --- | --- | --- | --- | --- | --- | --- | --- | --- | --- | --- | --- |
|  | Probe<br>Mis-<br>match<br>locatio<br>n |  |  |  | M1-F<br>Mis-<br>match<br>location |  |  |  | M1-R<br>Mis-<br>match<br>locatio<br>n |  |  |  |
|  | Intern<br>al | 3<br>' | centr<br>al | 5' | Internal | 3<br>' | centr<br>al | 5<br>' | intern<br>al | 3' | centra<br>l | 5' |
| **H1N1** |  |  |  |  |  |  |  |  |  |  |  |  |
| LC662543.1 | All | 1 | 0 | 0 | 0 | 0 | 0 | 0 | 0 | 0 | 0 | 0 |
| NC_002016 | All | 1 | 0 | 0 | 0 | 0 | 0 | 0 | 0 | 0 | 0 | 0 |
| MK680249.1 | All | 1 | 0 | 0 | 0 | 0 | 0 | 0 | 0 | 0 | 0 | 0 |
| **H3N2** |  |  |  |  |  |  |  |  |  |  |  |  |
| KJ609209.1 | All | 3 | 3 | 0 | 0 | 0 | 0 | 0 | 0 | 0 | 0 | 0 |
| NC_007367.1 | All | 3 | 3 | 0 | 0 | 0 | 0 | 0 | 0 | 0 | 0 | 0 |
| PQ731530.1 | All | 3 | 3 | 0 | 0 | 0 | 0 | 0 | 0 | 0 | 0 | 0 |
| **H5N1** |  |  |  |  |  |  |  |  |  |  |  |  |
| KY614875.1 | All | 1 | 0 | 1 | 0 | 0 | 0 | 0 | All | 0 | 0 | 2 |
| PP599468.1 | All | 1 | 0 | 0 | All | 0 | 0 | 1 | 0 | 0 | 0 | 0 |
| OR8193403.1 | All | 1 | 0 | 0 | All | 0 | 0 | 1 | 0 | 0 | 0 | 0 |

*Central (positions 7–13) → most impact on fluorescence. Terminal (positions 1–3 or* *17–19) → least impact*

The positional weighting analysis indicated that the majority of mismatches occurred outside the central probe region (positions 7–13), which is generally considered most critical for probe hybridization efficiency (Table 3b). Thus, **the assay is mismatch-tolerant because despite the presence of mismatches, the assay remains theoretically functional because mismatches occur at positions known to be thermodynamically and kinetically tolerated** (Table 3b).

### Quantitative mismatch analysis

Primer and probe mismatches were enumerated and quantified by aligning oligonucleotide binding regions against all 28 sequences and mapping substitutions to specific nucleotide positions. No mismatches were detected in primer regions among H1N1 and H3N2 sequences. Probe mismatches were present in all H3N2 sequences (3/3) and in three of four H5N1 sequences (Table 4)

**Table 4:**
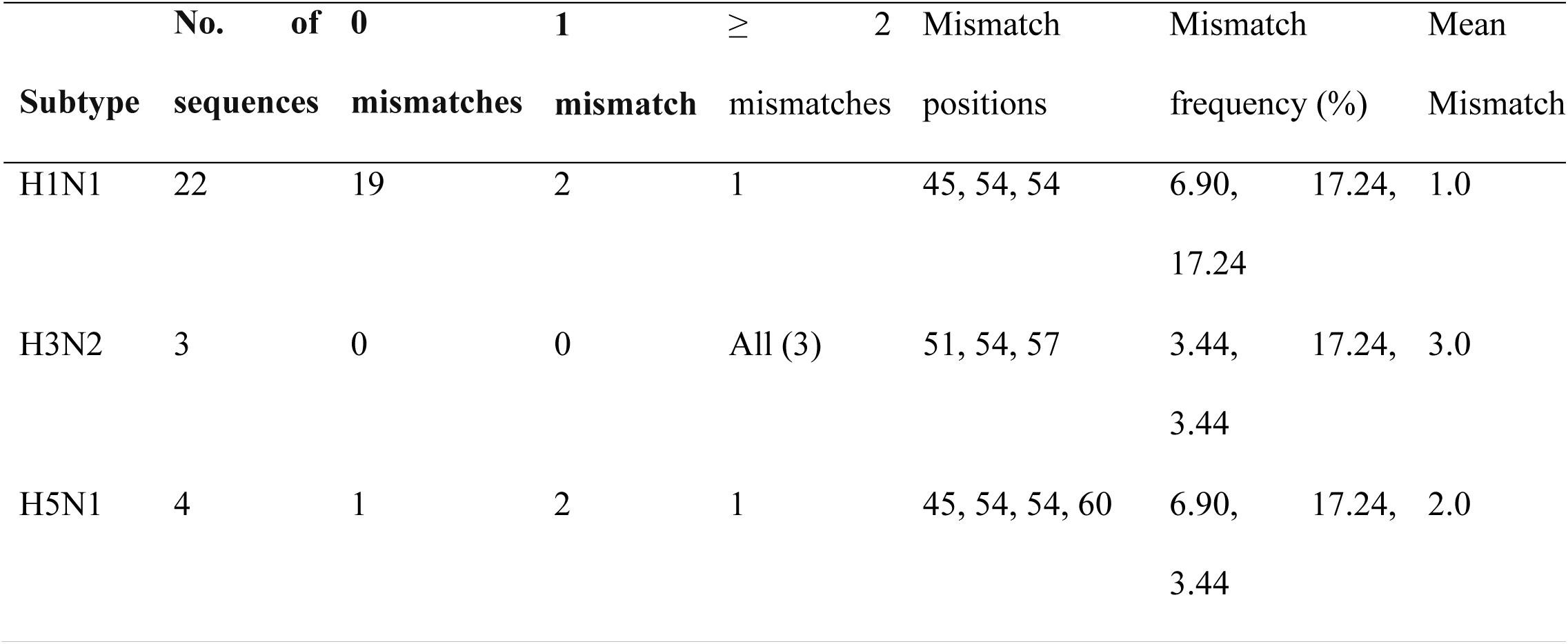
Mismatch quantification Table.

The mean number of probe mismatches per sequence was lowest in H1N1 (mean = 1.0), intermediate in H3N2 (mean = 3.0), and highest in H5N1 (mean = 2.0). Across all non-H1N1 isolates, the mean mismatch was 2.43 per isolate. Most mismatches occurred at internal probe positions, with a single terminal 3′ probe mismatch observed only in the historical H5N1 isolate (Table 4).

### Probe-region variation heatmap

A binary heatmap was constructed to visualize mismatch distribution across the 19-bp probe region (positions 42–60) relative to the H1N1 consensus sequence (Figure 3). H1N1 sequences showed complete probe conservation. H3N2 and H5N1 sequences exhibited subtype-specific internal mismatches, while only the historical H5N1 isolate demonstrated a terminal 3′ mismatch. Recent H5N1 isolates displayed a conserved internal mismatch pattern, suggesting stability of probe variation in contemporary lineages.

**Figure 3:**
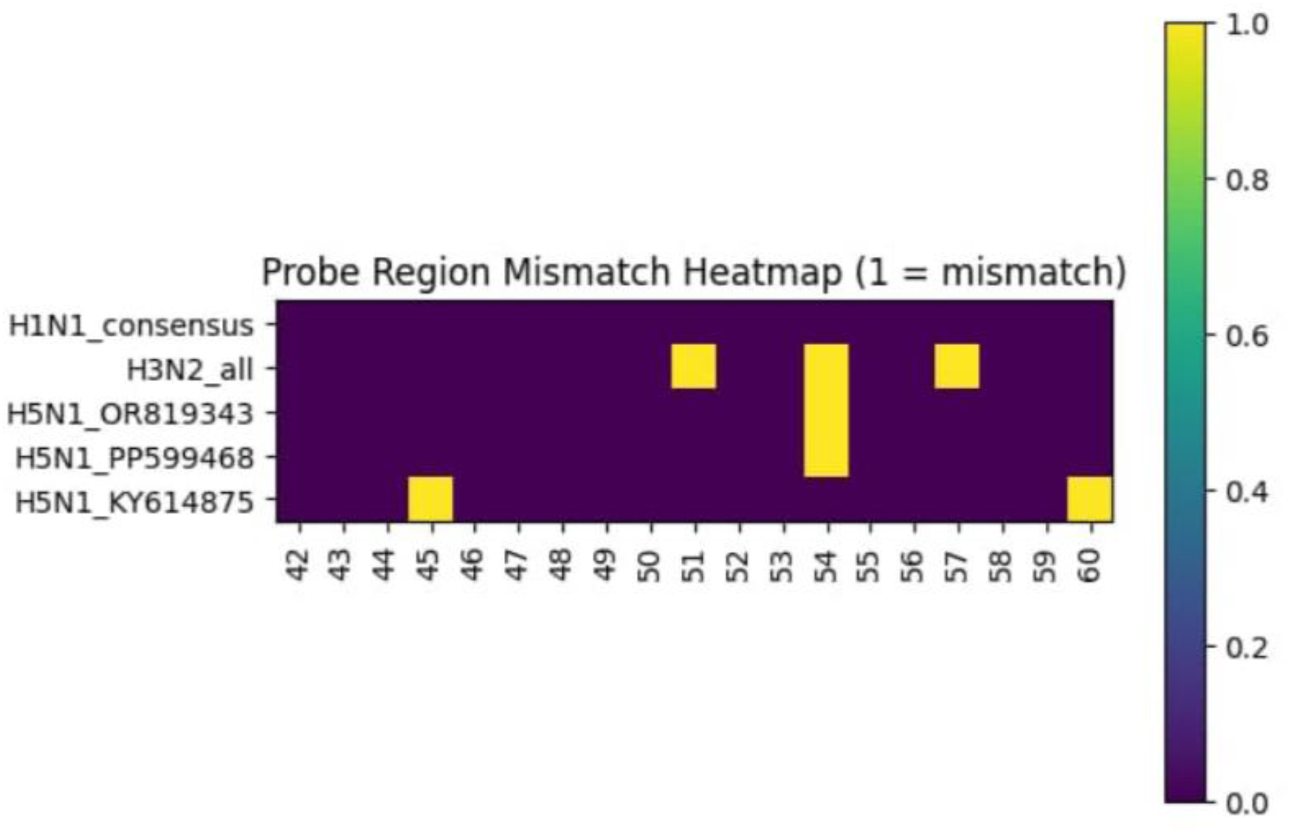
Probe-region variation heatmap.

### Estimated Ct Impact of Probe Mismatches

The potential impact of probe mismatches on cycle threshold (Ct) values was estimated based on previously published real-time PCR performance data [9]. Expected effects were categorized as follows: **Perfect match:** baseline Ct; **One internal probe mismatch:** +0.3 to +1.0 Ct; **Two internal probe mismatches:** +0.8 to +1.5 Ct; **Three or more internal probe mismatches:** potential loss of assay sensitivity. These estimates were used solely for theoretical interpretation of mismatch tolerance and were not intended to replace experimental validation.

### In silico specificity analysis using Primer-BLAST

Primer specificity was evaluated using NCBI Primer-BLAST against the nucleotide collection (nt) database. When no organism restriction was applied, no unintended amplification products were detected outside Influenza A virus sequences (Figure 4a). Restricting the analysis to Homo sapiens, Sus scrofa, and Canis lupus familiaris yielded no predicted amplification products, indicating absence of cross-reactivity with host genomes (Figure 4b).

**Figure 4a:**
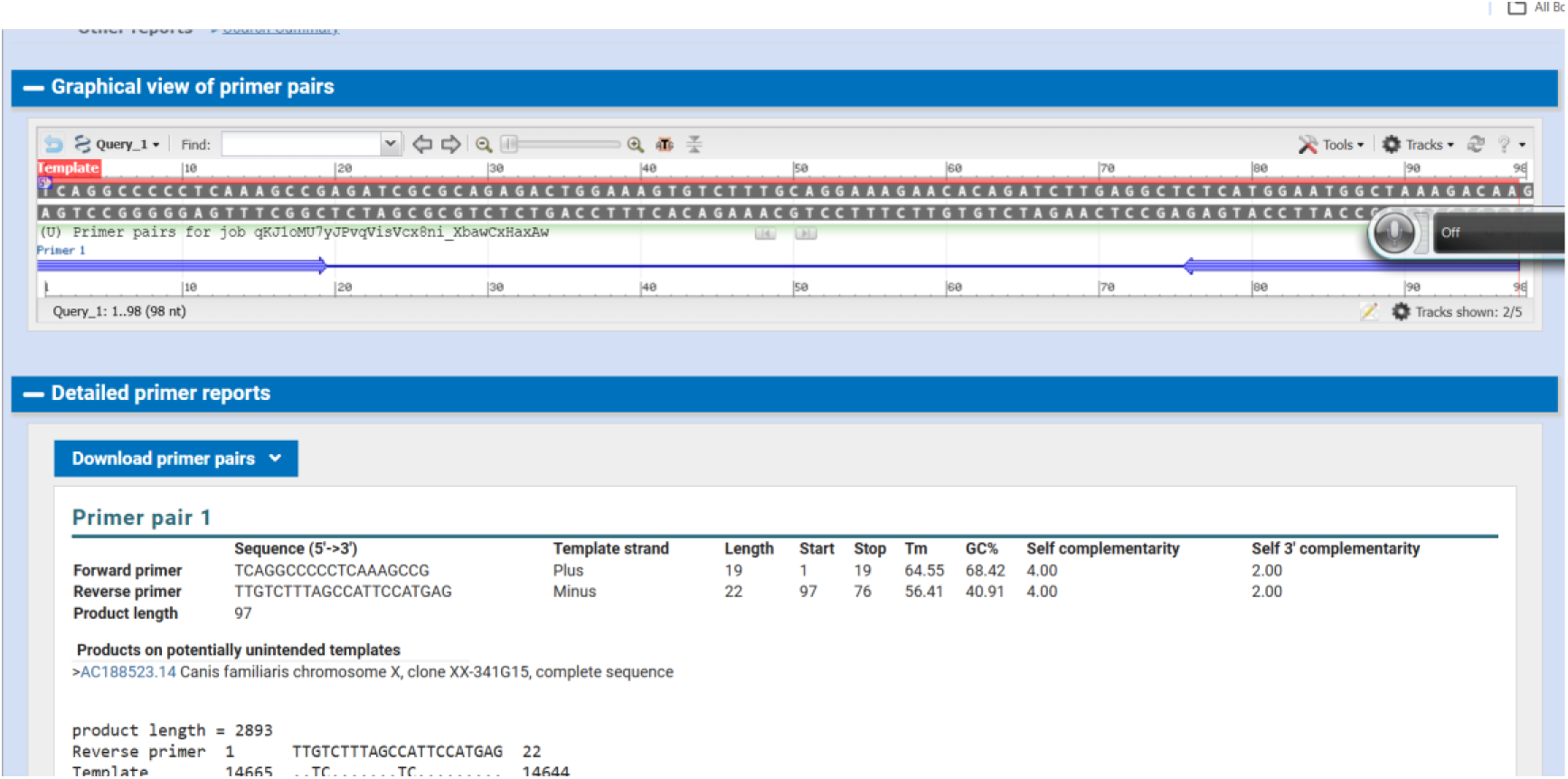
Primer Blast using blank.

**Figure 4b:**
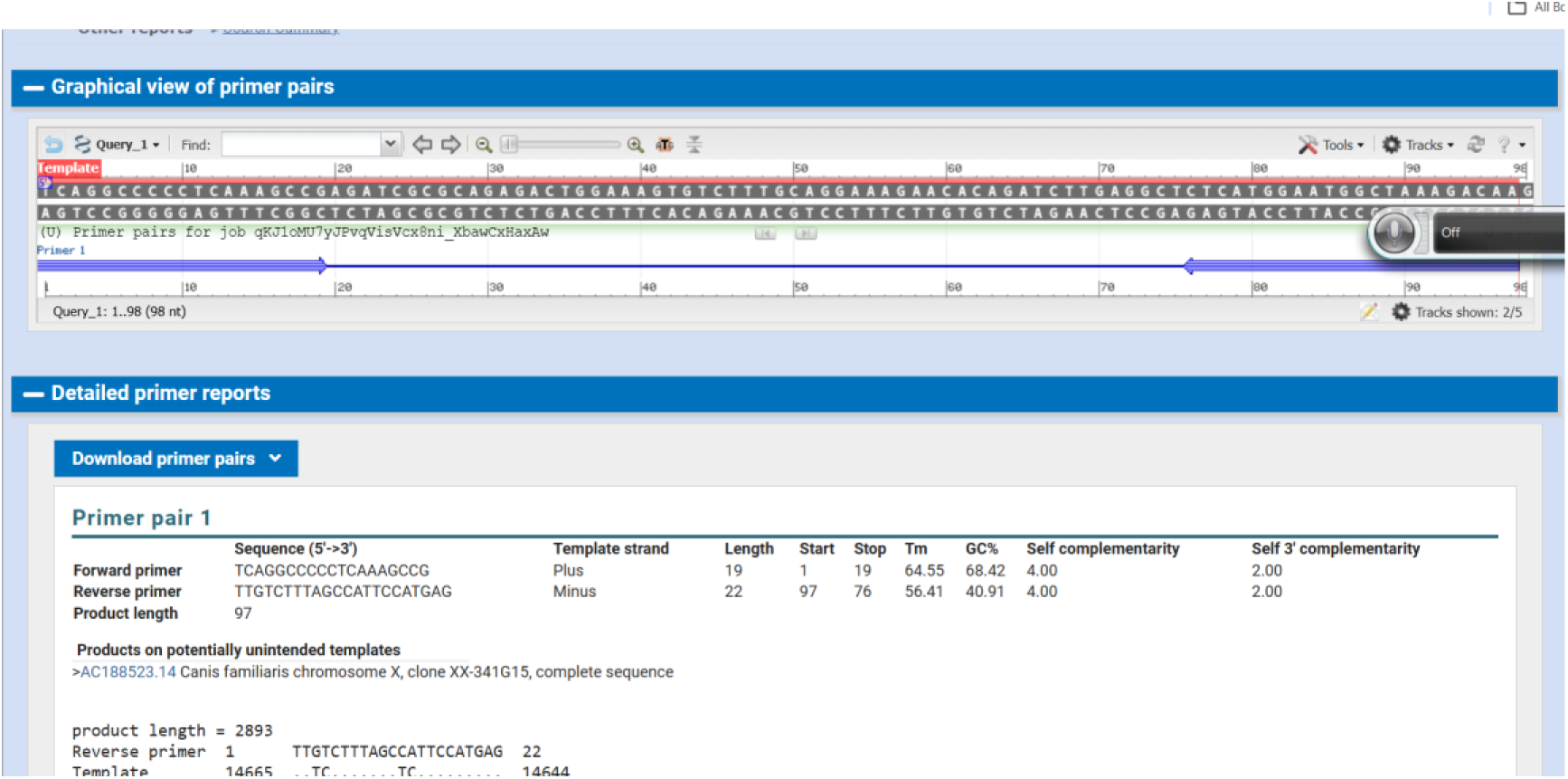
Primer-BLAST was restricted to Homo sapiens, Sus scrofa, and Canis lupus familiaris,

**Figure 4c:**
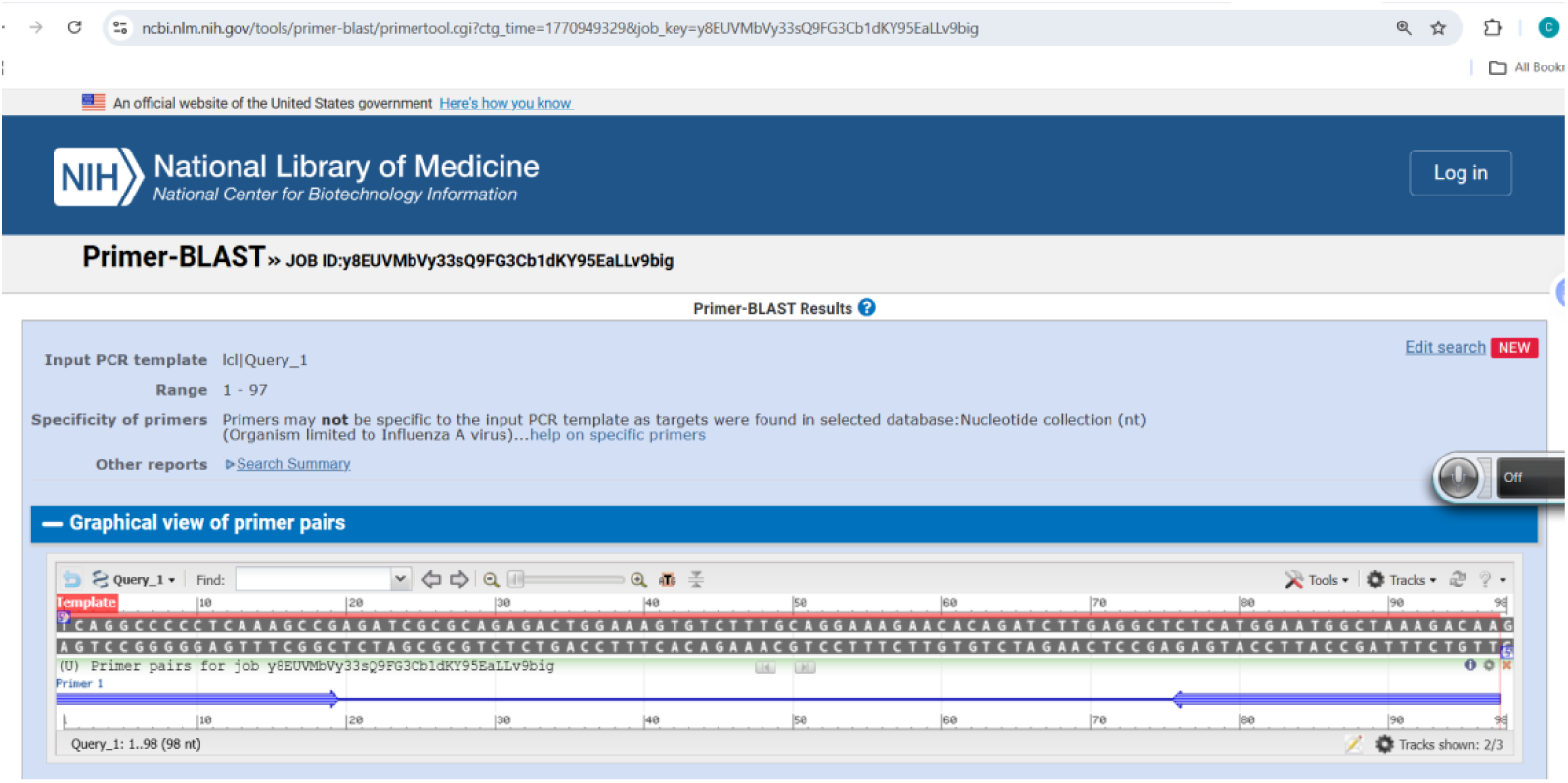
Primer Blast with organism restricted to influenza A virus.

**Figure 4d:**
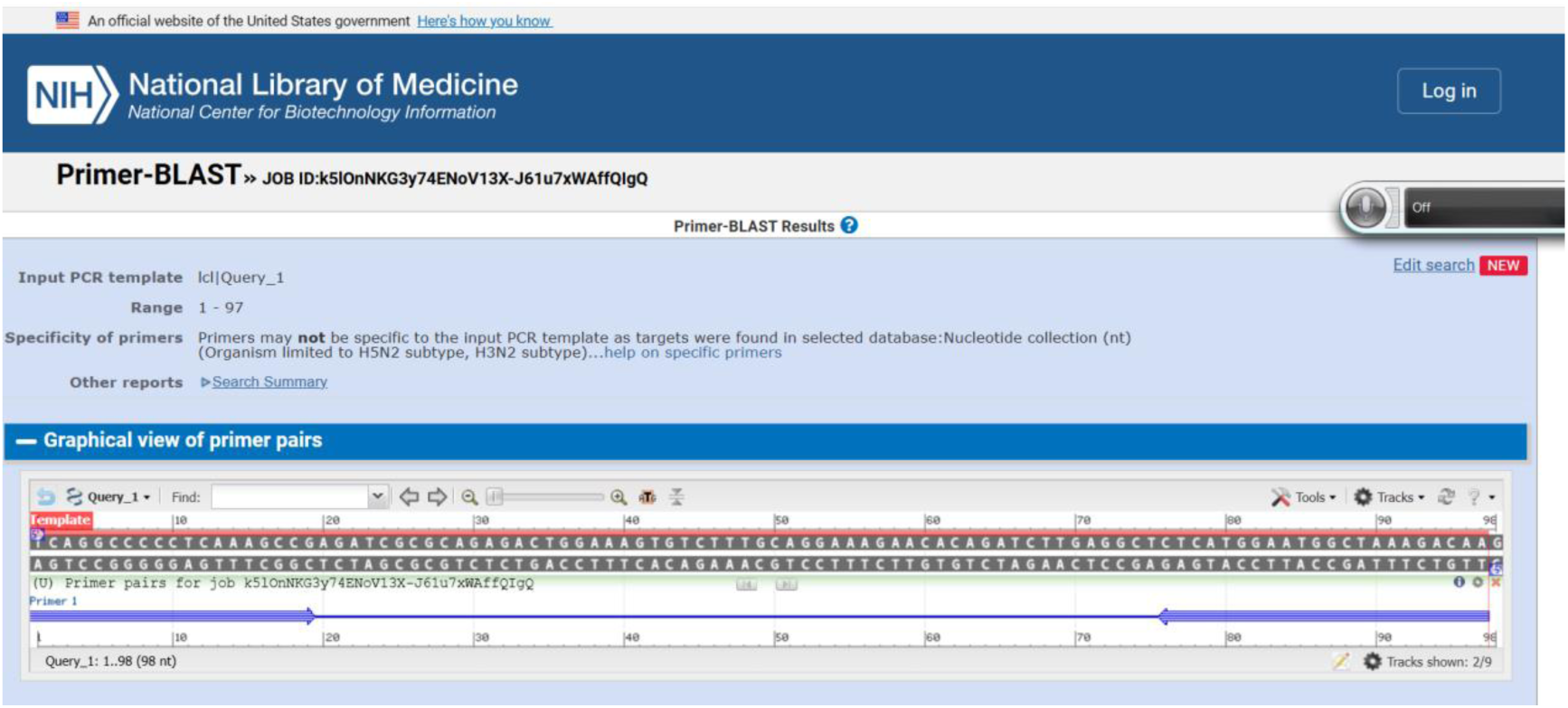
Primer Blast result with organism restricted to H1N1, H3N2 and H5N1 subtype.

When Primer-BLAST was restricted to Influenza A virus sequences, the primer pair consistently generated a predicted amplicon of the expected size (∼97 bp) (Figure 4c). Additional analyses restricted to H1N1, H3N2, and H5N1 subtypes yielded similar amplification profiles, indicating broad subtype coverage (Figure 4d).

### Assessment of primer–dimer formation

Computational analysis of self-complementarity and 3′ complementarity revealed low predicted primer–dimer potential for both primers. No high-risk self- or heterodimer interactions were identified, indicating a low likelihood of primer–dimer formation under standard amplification conditions.

### Interpretation of mismatch tolerance

Although nucleotide mismatches were observed within the probe region of H3N2 and H5N1 sequences, these were predominantly internal and spared primer 3′ termini. A single terminal probe mismatch was observed in one historical H5N1 isolate. Based on published RT-qPCR thermodynamic principles, internal probe mismatches are expected to primarily reduce hybridization efficiency rather than prevent amplification, whereas primer 3′ mismatches are more likely to impair extension. The observed mismatch patterns are therefore consistent with theoretical assay tolerance to subtype-associated sequence variation.

## Discussion

Genetic drift and host-associated adaptation continue to challenge the long-term reliability of molecular diagnostics for influenza A virus detection. RT-qPCR assays, while highly sensitive and specific, are intrinsically vulnerable to nucleotide mismatches within primer and probe binding regions, which can reduce amplification efficiency or lead to false-negative results. This risk is amplified in surveillance settings where assays are applied across multiple influenza A subtypes and hosts [3–4]. In this study, we performed a systematic in silico evaluation of primer and probe sequence variability within a matrix (M) gene–targeted RT-qPCR assay, with particular emphasis on mismatch tolerance across contemporary human and zoonotic strains.

Alignment of 22 H1N1 isolates with representative H3N2 and H5N1 reference sequences revealed conservation of primer and probe binding regions within H1N1, confirming the suitability of the assay for its primary target. In contrast, non-H1N1 subtypes exhibited varying degrees of sequence divergence, predominantly within the probe region. Across seven non-H1N1 reference isolates, a total of eleven mismatches were identified, yielding a mean mismatch of 2.43 per isolate. Importantly, these mismatches were unevenly distributed across assay components, with probe mismatches accounting for the majority of observed variability.

All three H3N2 isolates demonstrated probe mismatches, each involving 3 single internal nucleotide substitution, while no primer mismatches were detected. This finding is consistent with prior reports that internal probe mismatches may increase cycle threshold (Ct) values without necessarily abolishing detection, particularly when primer binding remains fully conserved [6]. Such probe-only mismatch profiles suggest that amplification efficiency is preserved, while fluorescence kinetics may be modestly affected. The uniformity of mismatch patterns across H3N2 isolates further indicates subtype-specific evolutionary signatures within the M gene probe target.

Greater heterogeneity was observed among H5N1 isolates. Two recent zoonotic isolates (OR819343.1 and PP599468.1), obtained in 2023 and 2024 from avian and bovine hosts respectively, shared identical A→G substitutions in both the probe region and the forward primer. Notably, these mismatches were internal and did not involve primer 3′ terminal nucleotides, which are known to be critical for polymerase extension. The recurrence of identical substitutions in temporally and epidemiologically distinct isolates suggests selective tolerance of these variants within circulating H5N1 lineages, while remaining compatible with assay detection.

A third H5N1 isolate (KY614875.1), collected in 2016 from a chicken host in Indonesia, displayed a more complex mismatch profile, including two probe mismatches, one internal and one at the probe’s 3′ terminal nucleotide, as well as two reverse primer mismatches. Although terminal probe mismatches are traditionally considered higher risk than internal substitutions, probes do not serve as substrates for polymerase extension, and their performance is primarily governed by hybridization stability and cleavage efficiency. Consequently, isolated terminal probe mismatches may reduce signal intensity or delay Ct values without necessarily preventing detection [6,10]. The absence of this mismatch pattern in more recent H5N1 isolates further suggests that this configuration represents a historical lineage rather than a dominant contemporary threat to assay performance. By implication, the observed mismatch patterns are consistent with distance-dependent genetic divergence previously described for influenza surface glycoproteins such as neuraminidase [11], although occurring at a reduced rate due to stronger functional constraint on the matrix gene.

Collectively, these findings underscore a key principle in molecular diagnostics: **the functional impact of mismatches depends on their number, position, and context, rather than their mere presence [10].** Primer mismatches were infrequent and largely avoided 3′ terminal positions, supporting sustained amplification across subtypes. Probe mismatches, while more common, were predominantly internal and occurred in patterns consistent with partial tolerance. These characteristics indicate that the evaluated M gene RT-qPCR assay exhibits inherent mismatch resilience, enabling detection across divergent influenza A subtypes and host species.

From a surveillance and pandemic preparedness perspective, this mismatch tolerance is particularly valuable. Influenza A viruses circulate at the human–animal interface and are subject to frequent reassortment and host-driven selection pressures [1–2, 12]. Diagnostic assays that are overly dependent on perfect sequence identity risk silent failure as viral genomes evolve [7,13–14]. By contrast, assays designed and evaluated with mismatch resilience in mind are better suited for long-term deployment in both clinical and One Health surveillance contexts [15].

This study is limited by its in silico design, and experimental validation is required to quantify the precise effects of observed mismatches on analytical sensitivity and Ct shifts. Future work incorporating synthetic templates or viral RNA representing high-risk mismatch profiles will be essential to confirm functional tolerance. Nonetheless, the present analysis provides a practical framework for integrating mismatch mapping into routine assay evaluation and highlights the importance of proactive diagnostic resilience assessment in the face of viral evolution.

## Acknowledgement

We sincerely appreciate ChatGPT and its manufacturers for mentoring me through this project.

## Conflict of interest

There is no conflict of interest Ethical consideration

The study was carried out in line with ethics guiding bioinformatics studies

## Availability of data

Data used in this study were obtained from public data base (NCBI).

